# MRI-Based Glymphatic Markers and Their Associations with Polysomnographic Sleep in Relapsing-Remitting Multiple Sclerosis

**DOI:** 10.64898/2026.09.30.755605

**Authors:** Roya Sharifpour, Sophie Elands, Jeromy Hrabovecky, Antonin Rovai, Serena Borrelli, Gabrielle de Bourgoing, Gaetano Perrotta, Xavier De Tiège, Mélanie Strauss

**Author notes:** These authors contributed equally to this work (Roya Sharifpour, Sophie Elands). Corresponding author: Mélanie Strauss Université libre de Bruxelles (ULB), Hôpital Universitaire de Bruxelles (H.U.B), CUB Hôpital Érasme, Department of Neurology, Brussels, Belgium.

## Abstract

**Background and Objectives:** The glymphatic system is the brain’s waste clearance pathway. Although its dysfunction has been associated with several neurological disorders, its role in multiple sclerosis (MS) remains poorly understood. Glymphatic activity is thought to peak during sleep, yet its relationship with objectively measured sleep has scarcely been explored in humans. We assessed two MRI glymphatic proxies in patients with relapsing-remitting MS (PwMS) and healthy controls (HCs) and their associations with polysomnographic measures, structural MRI, and, in PwMS, disease burden.

**Methods:** In this cross-sectional study, PwMS (Expanded Disability Status Scale [EDSS] ≤4.0) and matched HCs underwent overnight polysomnography and 3T MRI. Glymphatic proxies were the diffusion tensor image analysis along the perivascular space (DTI-ALPS) index and coupling of the global blood-oxygen-level–dependent signal with CSF inflow (gBOLD–CSF coupling). We also assessed sleep micro- and macroarchitecture, EDSS, disease duration, lesion volume, mean diffusivity (MD), lateral ventricle (LV) and choroid plexus volumes. Analyses were adjusted for age and sex.

**Results:** We included 22 PwMS (34.2±10.1 years; 16 female) and 26 HCs (32.2±5.7 years; 18 female). DTI-ALPS was lower in PwMS (adjusted difference: −0.12, 95% CI −0.22 to −0.02), but MD was higher, and adjustment for MD attenuated this difference (−0.11, −0.21 to −0.0002). gBOLD–CSF coupling did not differ between groups (−0.02, −0.08 to 0.05). In PwMS, stronger coupling was associated with shorter total sleep time (r=0.49, 95% CI 0.05 to 0.77) and higher arousal index (r=−0.48, −0.77 to −0.04), whereas DTI-ALPS was unrelated to sleep. Associations with clinical and structural measures were borderline for DTI-ALPS and EDSS (r=−0.44, −0.74 to 0.01) and for coupling and LV volume (r=0.44, −0.00 to 0.74), and weak otherwise. Analyses were exploratory and uncorrected for multiplicity.

**Discussion:** In MS, the DTI-ALPS index is partly confounded by white matter damage and should not serve as a standalone glymphatic marker. Stronger gBOLD– CSF coupling was associated with shorter, fragmented sleep in PwMS, consistent with the proposed compensatory waking CSF dynamics. If confirmed longitudinally, gBOLD–CSF coupling may serve as a glymphatic-related marker in MS and sleep a potential target to improve brain clearance in PwMS.

## Introduction

Multiple sclerosis (MS) is a chronic immune-mediated disease of the central nervous system (CNS) characterized by inflammation, demyelination, and neurodegeneration that lead to cumulative disability.^1^ Lesions develop around small veins and postcapillary venules, where peripheral T and B cells that have crossed a disrupted blood–brain barrier accumulate in the perivascular spaces and release soluble factors that induce demyelination, directly or through microglial activation.^2^ Although MS is initiated by autoimmune mechanisms, the persistence and spatial spread of inflammation may also depend on how efficiently the CNS removes proinflammatory cytokines, myelin debris, and other solutes from the interstitial fluid.^3^

The glymphatic system is a brain-wide perivascular network that supports this clearance. CSF enters periarterial spaces, exchanges with interstitial fluid through astrocytic aquaporin-4 (AQP4) channels, and drains along perivenous routes toward the lymphatic system.^4,5^ Glymphatic transport depends on perivascular space integrity, arterial pulsatility, and AQP4-mediated exchange ^4,5^, each of which may be altered in MS.^6^ Conversely, impaired clearance could prolong exposure to inflammatory mediators and promote secondary neurodegeneration, suggesting a bidirectional relationship between glymphatic dysfunction and neuroinflammation.

Glymphatic activity is strongly regulated by the sleep–wake cycle and peaks during non–rapid eye movement (NREM) sleep.^7^ Sleep disturbance is common in MS and includes reduced total sleep time and sleep efficiency, increased fragmentation, and altered NREM and REM sleep architecture.^8^ Sleep disturbance is therefore a plausible modifier of glymphatic function in MS, yet the relationship between objective sleep measures and imaging markers of glymphatic activity has received little attention in this disease, and current available evidence relies on sleep questionnaires rather than polysomnography (PSG).^9^

Contrast-enhanced MRI after intrathecal gadolinium remains the reference method for assessing glymphatic function in humans,^10^ but it is invasive and carries potential adverse effects. Non-invasive proxies have therefore been developed.^11^ The most widely used, the diffusion tensor image analysis along the perivascular space (DTI-ALPS) index, quantifies water diffusivity along the perivenous spaces of medullary veins at the level of the lateral ventricle bodies.^12^ A lower index has been reported in many neurological diseases, including MS, and is commonly interpreted as glymphatic impairment.^3,11,13^ This interpretation has been increasingly criticized,^14,15^ and the developers of the method have cautioned that a lower index primarily reflects reduced left–right (x-axis) diffusivity in periventricular white matter (WM) rather than glymphatic dysfunction, acknowledging that the title of their original report ^12^ may have overstated this link.^16^ Because the index is derived from WM diffusivity, it may be particularly susceptible to the WM microstructural damage that characterizes MS, a potential confound that has rarely been addressed.

Resting-state fMRI is another approach that has gained interest. Low-frequency fluctuations of the global blood-oxygen-level–dependent (gBOLD) signal are temporally coupled with cerebrospinal fluid (CSF) inflow, a phenomenon first described during sleep^17^ and also observed during wakefulness, and this gBOLD– CSF coupling is thought to index glymphatic-related fluid dynamics.^18,19^ Global hemodynamic fluctuations, which are closely linked to arousal-related changes in neural activity, alter cerebral blood volume and because intracranial volume is fixed (Monro–Kellie doctrine), thereby drive CSF movement into and out of the brain, providing a physiologic link between brain activity and CSF dynamics.^17,19^ Weaker coupling has been reported in Alzheimer disease, Parkinson disease, and chronic insomnia,^18,20,21^ but, to our knowledge, it has not been examined in MS.

Choroid plexus (ChP) and lateral ventricle (LV) enlargement are frequent in MS and may influence CSF dynamics.^22,23^ ChP enlargement has been associated with reduced CSF secretion in Alzheimer disease,^24^ whereas LV enlargement mainly reflects atrophy,^23^ which is accompanied by perivascular space enlargement in MS,^6^ and ventricular volume changes may also mechanically drive CSF flow.^25^

In this study, we combined DTI-ALPS and gBOLD–CSF coupling with overnight PSG in patients with relapsing-remitting MS (PwMS) and healthy controls (HCs). We additionally measured mean diffusivity (MD) within the DTI-ALPS regions to quantify the contribution of local WM microstructure to the index, and related both proxies to disability, disease duration, lesion volume, and ChP and LV volumes. We hypothesized that (1) glymphatic proxies would be reduced in patients; (2) in patients, these proxies would be associated with sleep measures, disability, disease duration, lesion volume, and CSF-compartment volumes; and (3) WM microstructural damage would confound the DTI-ALPS index.

## Methods

The study was approved by the Ethics Committee of Hôpital Universitaire de Bruxelles and Université libre de Bruxelles (reference SRB2020_394) and conducted in accordance with the Declaration of Helsinki. All participants provided written informed consent.

### Participants

Between September 2021 and December 2024, we enrolled 26 PwMS and 28 HCs aged 18–50 years (mean±SD: 33.0±8.0). Patients were recruited from the MS outpatient clinics of Hôpital Érasme. HCs were matched to patients for age, sex, and education. Patients had clinically definite relapsing-remitting MS (RRMS) according to the 2017 McDonald criteria,^26^ an Expanded Disability Status Scale (EDSS) score ≤4.0,^27^ and no relapse or corticosteroid treatment within the preceding 6 weeks. Exclusion criteria for all participants were another major neurological or psychiatric disorder, an untreated sleep disorder, body mass index >30, pregnancy or breastfeeding, alcohol, benzodiazepine, or other substance misuse, and contraindications to MRI.

### Clinical and Sleep Assessments

EDSS, disease duration, and disease-modifying therapy (**Supplementary Table S1**) were obtained by a neurologist (S.E.). Handedness was assessed with the Edinburgh Handedness Inventory.^28^ Overnight PSG was performed in the hospital or as an ambulatory recording (Morpheus Recorder, OSG BV, Kontich, Belgium) with participants sleeping in a nearby hotel, using EEG electrodes positioned according to the international 10–20 system. Sleep stages and arousals were scored manually in BrainRT (OSG BV) according to American Academy of Sleep Medicine criteria^29^ by scorers blinded to group. Macrostructural measures were total sleep time (TST) and arousal index, the latter reflecting sleep fragmentation. Because glymphatic activity peaks during deep sleep, slow-wave activity (SWA) and slow-wave density (SWD) during N3 were computed with the YASA toolbox^30^ as microstructural measures.

### MRI Acquisition

All participants were scanned on the same 3T SIGNA PET/MR system (GE Healthcare, Chicago, IL) the morning after PSG between 9 and 11 AM to limit time-of-day effects on perivascular measures. The protocol included 3D fluid-attenuated inversion recovery (FLAIR), 3D T1-weighted inversion-recovery spoiled gradient-recalled echo, multishell diffusion-weighted echo-planar imaging (DWI), 3D susceptibility-weighted imaging (SWI), and resting-state gradient-echo echo-planar functional MRI (rs-fMRI) during which participants were instructed to fixate on a cross. Acquisition parameters are provided in the Supplementary Methods.

### Lesion Segmentation and Volumetry

T2-FLAIR hyperintense lesions were segmented with LST-AI^31^ and verified and manually corrected by an experienced MS neurologist (S.E.). Lesion filling was performed on T1-weighted and FLAIR images before volumetric analyses to limit the effect of lesions on automated segmentation. LV volumes were derived from lesion-filled T1-weighted images with FreeSurfer 6.0.1,^32^ and ChP volumes with the method of Tadayon et al.,^33^ which outperforms the FreeSurfer ChP segmentation. All segmentations were reviewed by S.E., blinded to clinical data, and normalized to estimated total intracranial volume (eTIV) (**Supplementary Methods**).

### DTI-ALPS Index

DWI and SWI preprocessing are described in the Supplementary Methods. Mean MD and diffusivity maps along the x-, y-, and z-axes (Dxx, Dyy, Dzz) were derived from the fitted tensor. SWI images and lesion masks were coregistered to DWI space, and the axial slice at the level of the lateral ventricle bodies in which medullary veins run perpendicular to the ventricular wall was selected (**Figure 1**). Because the DTI-ALPS index is derived from WM regions involved in motor and language function and varies with handedness,^34^ it has been recommended to compute the index in the right hemisphere when handedness is unknown, or in the left hemisphere of right-handed participants. Handedness was available for all participants; the 2 left-handed participants (1 PwMS, 1 HC) were therefore excluded to obtain a homogeneous sample, and the index was computed in the left hemisphere. On the color-coded fractional anisotropy map, S.E. placed spherical regions of interest (ROIs; radius 2 mm) in the projection and association fiber areas at least 2 voxels from lesion edges. In these regions, the principal fiber directions are along the z-axis (projection fibers) and the y-axis (association fibers), whereas medullary veins and their perivascular spaces run along the x-axis. The DTI-ALPS index was therefore calculated as the mean diffusivity parallel to the veins and perpendicular to the fibers (Dxx in the projection and association ROIs) divided by the mean diffusivity perpendicular to both (Dyy in the projection ROI and Dzz in the association ROI).^12^ All coregistrations and ROI placements were visually checked. MD averaged across both ROIs served as an index of local WM microstructural integrity.

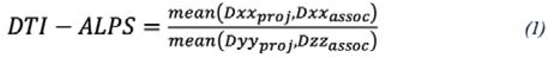

**Figure 1.**
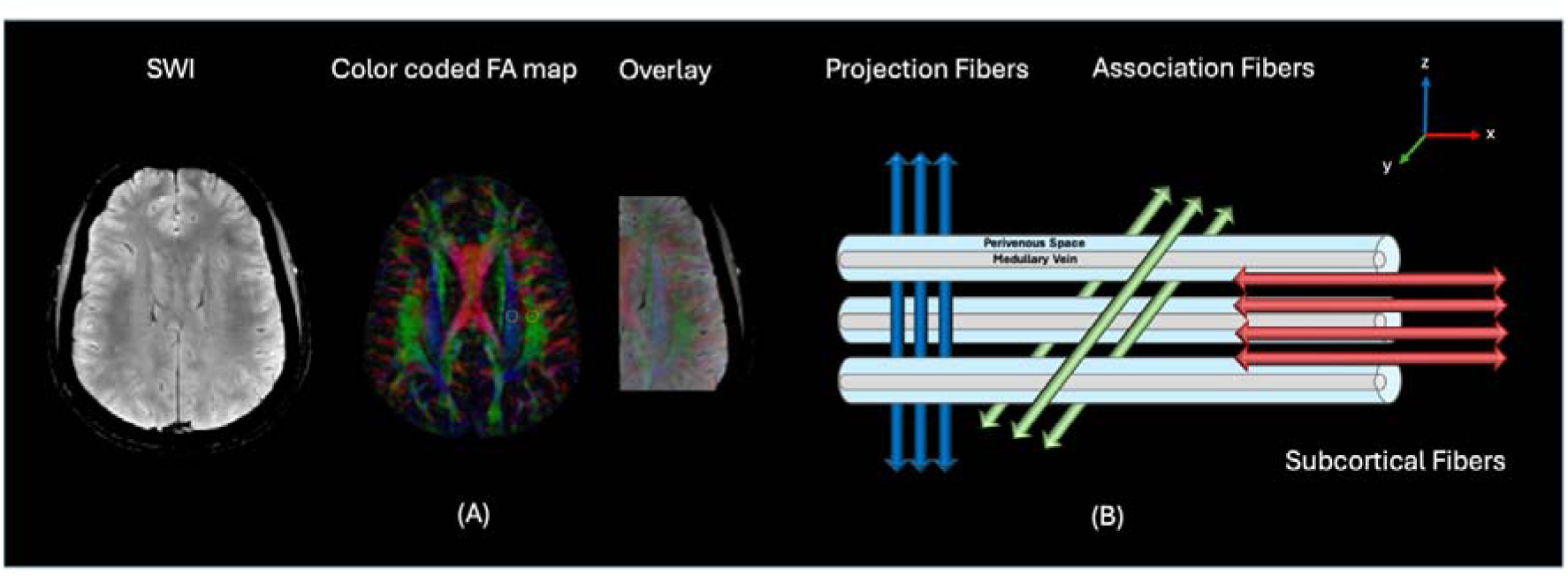
DTI-ALPS Calculation in a Healthy Participant. (A) Susceptibility-weighted images coregistered to diffusion-weighted image space, together with the color-coded fractional anisotropy map, were used to place 2 spherical regions of interest (radius 2mm) in the left hemisphere, one in the projection fiber area and one in the association fiber area. (B) Diffusivities along the x-, y-, and z-axes were extracted from each region. The DTI-ALPS index was calculated as the ratio of diffusivities perpendicular to the fiber tracts and parallel to the medullary veins (Dxx in the projection and association areas) to diffusivities perpendicular to both the fiber tracts and the medullary veins (Dyy in the projection area and Dzz in the association area). DTI-ALPS = diffusion tensor image analysis along the perivascular space; FA = fractional anisotropy; SWI = susceptibility-weighted imaging.

### gBOLD–CSF Coupling

Resting-state fMRI preprocessing, including the rationale for not regressing out global, CSF, and motion signals, is described in the Supplementary Methods. The gBOLD signal was obtained by averaging z-normalized time series across a cortical gray matter mask, and the CSF inflow signal was extracted from CSF voxels in the lowest slice of the acquisition, where inflow effects are maximal,^17^ and expressed as percentage change from its temporal mean (**Supplementary Methods**). Cross-correlations between the gBOLD and CSF signals were computed at lags from −12 to +12 seconds in steps of 1 TR (3 seconds). Because the group-averaged cross-correlation functions of patients, HCs, and the combined cohort showed their most prominent (negative) peak at +3 seconds (**Figure 2.A**), the correlation at this lag was used as the coupling measure, with more negative values indicating stronger coupling. To further characterize the dynamic relationship between the two signals, an additional cross-correlation analysis was performed between the CSF signal and the negative temporal derivative of the gBOLD signal (**Figure 2.B**).^17,18^ The group-averaged coupling was tested against a null distribution generated by pairing gBOLD and CSF signals from different participants (10,000 permutations; **Supplementary Methods**).

**Figure 2.**
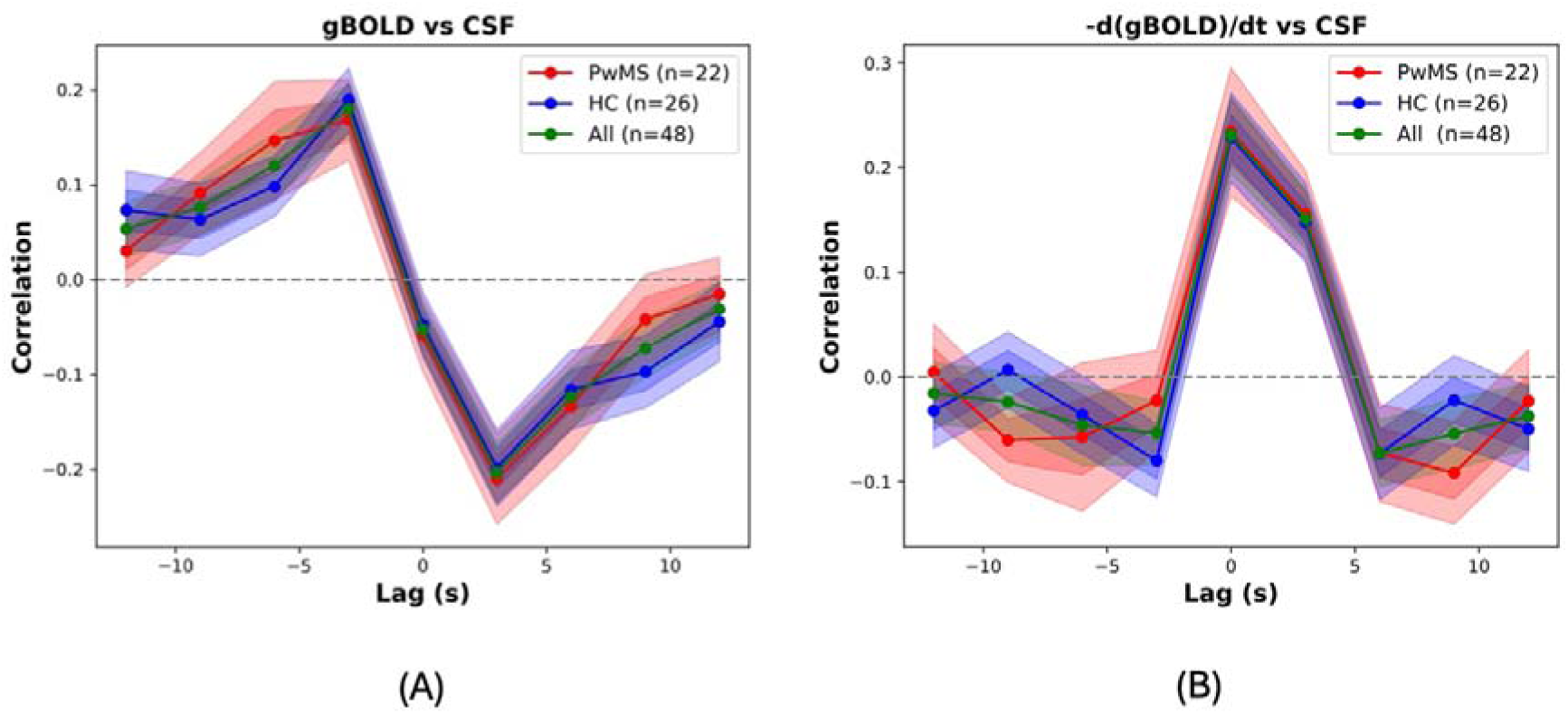
Coupling Between the Global BOLD Signal and CSF Inflow. (A) Mean cross-correlation between the gBOLD and CSF signals as a function of time lag. (B) Mean cross-correlation between the negative temporal derivative of the gBOLD signal (−d[gBOLD]/dt) and the CSF signal as a function of time lag. Results are shown for patients with relapsing-remitting multiple sclerosis (PwMS, red), healthy controls (HC, blue), and the combined cohort (All, green). Shaded areas indicate 95% CIs; the dashed line indicates zero correlation. gBOLD = global blood-oxygen-level–dependent signal.

### Statistical Analysis

The sample size was determined by recruitment during the study period. Within each group, values outside 1.5 times the interquartile range (IQR) were considered outliers; outlying participants (3 patients for DTI-ALPS, 1 HC for gBOLD–CSF coupling) were excluded from all analyses so that all analyses were performed in the same sample. Continuous variables were analyzed without categorization and are reported as mean ± SD, median (IQR), or median (range). Demographic differences are reported as mean differences, differences in proportions (Newcombe hybrid score interval), or Hodges–Lehmann median differences with 95% CIs. Group differences in glymphatic proxies, MD, structural MRI, and sleep measures were estimated with analysis of covariance (ANCOVA) adjusted for age and sex and are reported as adjusted mean differences (PwMS minus HCs) with 95% CIs; a secondary DTI-ALPS model was additionally adjusted for MD to estimate the contribution of local WM microstructure to the group difference. Associations between glymphatic proxies and sleep, clinical, and structural measures were assessed within each group with partial Pearson correlations adjusted for age and sex (and for MD in secondary DTI-ALPS models of clinical measures); associations with EDSS, disease duration, and lesion volume were assessed in PwMS only, with 95% CIs obtained by Fisher z transformation. The number of participants contributing to each analysis is given in **Tables 1 and 2**. Because the analyses were exploratory and not corrected for multiplicity, results are reported as effect estimates with 95% CIs and interpreted according to effect size and precision rather than statistical significance. Analyses were performed in Python 3.11 (SciPy, statsmodels, pingouin).

**Table 1.**
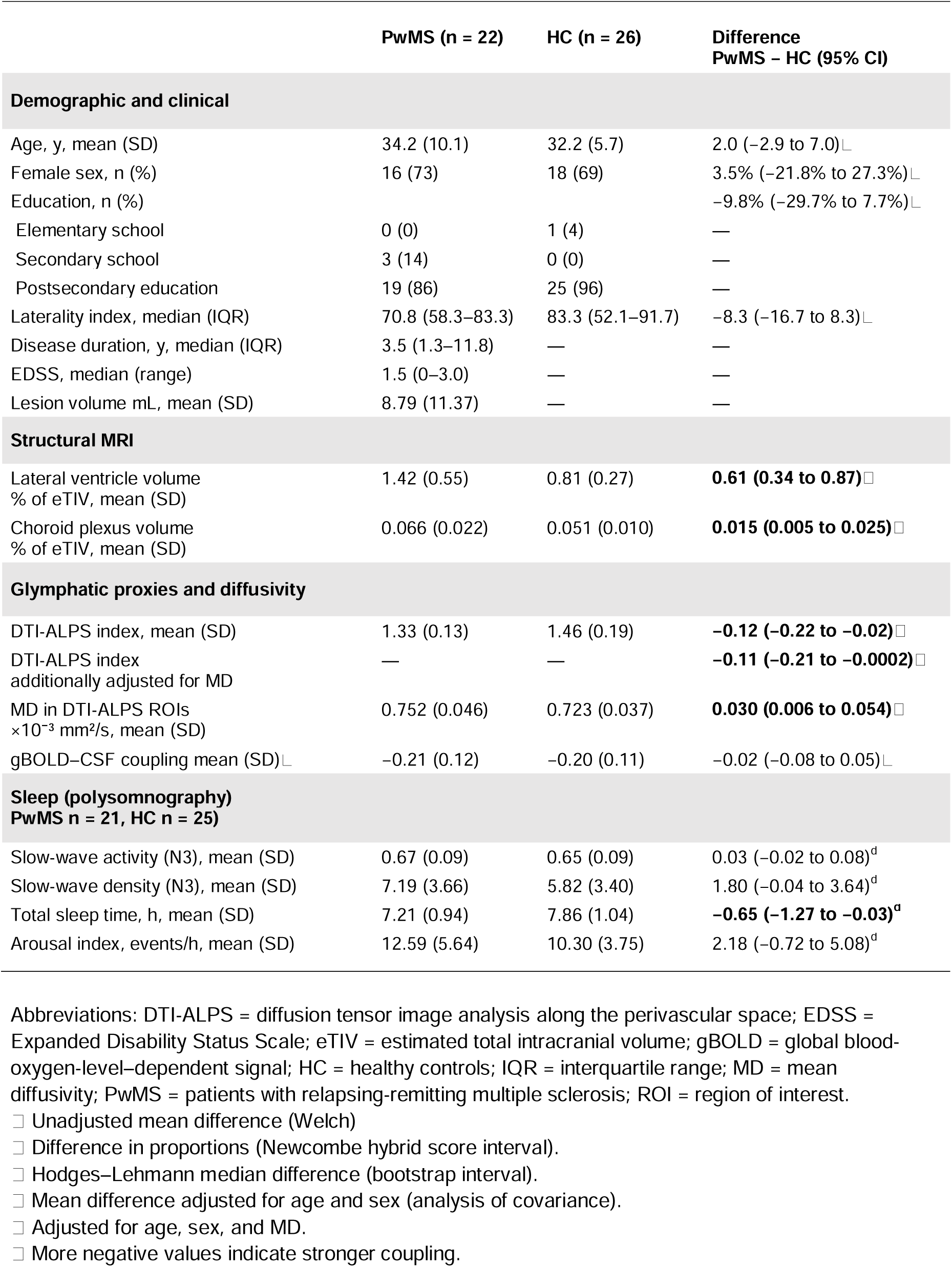
Demographic, Clinical, Structural MRI, Glymphatic, and Sleep Characteristics of the Participants.

**Table 2.** Partial Correlations of Glymphatic Proxies with Sleep, Structural MRI, Clinical, and Disease Burden Measures.

|  | DTI-ALPS |  | gBOLD–CSF Coupling |  |
|---|---|---|---|---|
|  | **HC**<br>r<br>(95% CI) | **PwMS**<br>r<br>(95% CI) | **HC**<br>r<br>(95% CI) | **PwMS**<br>r<br>(95% CI) |
| **Sleep microstructure**<br>**(HC: n = 25, PwMS: n = 21)** |  |  |  |  |
| Slow-wave activity (N3) | −0.06<br>(−0.46 to 0.36) | −0.05<br>(−0.49 to 0.41) | −0.03<br>(−0.44 to 0.39) | 0.17<br>(−0.30 to 0.58) |
| Slow-wave density (N3) | **−0.43**<br>**(−0.72 to −0.03)** | 0.05<br>(−0.41 to 0.49) | 0.36<br>(−0.06 to 0.67) | −0.16<br>(−0.57 to 0.32) |
| **Sleep macrostructure**<br>**(HC: n = 25, PwMS: n = 21)** |  |  |  |  |
| Total sleep time | 0.35<br>(−0.08 to 0.66) | 0.02<br>(−0.44 to 0.47) | −0.19<br>(−0.56 to 0.24) | **0.49**<br>**(0.05 to 0.77)** |
| Arousal index | −0.24<br>(−0.59 to 0.19) | 0.32<br>(−0.15 to 0.68) | −0.12<br>(−0.51 to 0.31) | **−0.48**<br>**(−0.77 to −0.04)** |
| **Structural MRI**<br>**(HC: n = 26, PwMS: n = 22)** |  |  |  |  |
| Choroid plexus volume | −0.25<br>(−0.59 to 0.17) | 0.00<br>(−0.44 to 0.44) | 0.19<br>(−0.23 to 0.55) | 0.33<br>(−0.13 to 0.68) |
| Lateral ventricle volume | 0.08<br>(−0.33 to 0.47) | −0.26<br>(−0.63 to 0.21) | 0.19<br>(−0.23 to 0.55) | **0.44**<br>**(−0.00 to 0.74)** |
| **Clinical disability and disease burden**<br>**(PwMS: n = 22)** |  |  |  |  |
| EDSS | — | −0.44<br>(−0.74 to 0.01) | — | 0.25<br>(−0.22 to 0.62) |
| EDSS,<br>additionally adjusted for MD | — | −0.43<br>(−0.74 to 0.03) | — | — |
| Disease duration | — | 0.23<br>(−0.23 to 0.61) | — | −0.22<br>(−0.60 to 0.25) |
| Disease duration,<br>additionally adjusted for MD | — | 0.28<br>(−0.20 to 0.65) | — | — |
| Lesion volume | — | −0.22<br>(−0.60 to 0.25) | — | 0.18<br>(−0.28 to 0.58) |

Values are partial Pearson correlation coefficients (95% CI) adjusted for age and sex. CIs were obtained with the Fisher z transformation. Because more negative gBOLD–CSF coupling values indicate stronger coupling, a positive r indicates weaker coupling with higher values of the measure. Analyses were exploratory and not corrected for multiple comparisons.
Abbreviations: DTI-ALPS = diffusion tensor image analysis along the perivascular space; EDSS = Expanded Disability Status Scale; gBOLD = global blood-oxygen-level–dependent signal; HC = healthy controls; MD = mean diffusivity; PwMS = patients with relapsing-remitting multiple sclerosis.

### Data Availability

Processed data and analysis code are available at https://github.com/ln2t/glymphatic-PwMS-paper. Requests for raw data may be addressed to the corresponding author and are subject to approval by the local ethics committee and a data transfer agreement.

## Results

### Demographic, Clinical and Sleep Characteristics

Of the 54 enrolled participants, 2 left-handed participants (1 PwMS, 1 HC), 3 PwMS with outlying DTI-ALPS values, and 1 HC with an outlying coupling value were excluded, leaving 22 PwMS and 26 HCs. PSG was unavailable for 1 PwMS and 1 HC (due to technical issues); sleep analyses therefore included 21 PwMS and 25 HCs. Groups were comparable in age, sex, education, and laterality index (**Table 1**). Patients had larger LV and ChP volumes and shorter TST than HCs, whereas differences in N3 SWA, N3 SWD, and arousal index were small, with CIs including zero (**Table 1**).

### Presence of gBOLD–CSF Coupling

In patients, HCs, and the combined cohort, group-averaged gBOLD–CSF cross-correlation functions showed a positive peak at −3 seconds (r = 0.17 to 0.19) and a negative peak at +3 seconds (r = −0.21 to −0.20; all P < 0.001 vs permutation null), and the CSF signal correlated with the negative temporal derivative of the gBOLD signal near zero lag (r = 0.23, P < 0.001; **Figure 2**). The negative correlation at +3 seconds indicates that increases in the gBOLD signal are followed by decreases in CSF inflow signal, and the near-zero-lag correlation with the negative gBOLD derivative indicates that CSF inflow increases when global blood volume decreases. This pattern replicates previous reports in healthy individuals and patient populations.^17,18^ The shape and amplitude of the cross-correlation functions were similar in PwMS and HCs.

#### Group Differences in Glymphatic Proxies

The DTI-ALPS index was lower in patients than in HCs (1.33±0.13 vs 1.46±0.19; adjusted difference −0.12, 95% CI −0.22 to −0.02). MD within the DTI-ALPS ROIs was higher in patients (0.752±0.046 vs 0.723±0.037 × 10 ³ mm²/s; adjusted difference 0.030, 95% CI 0.006 to 0.054), and additional adjustment for MD attenuated the group difference in DTI-ALPS (−0.11, 95% CI −0.21 to −0.0002). gBOLD–CSF coupling did not differ between groups (−0.21±0.12 vs −0.20±0.11; adjusted difference −0.02, 95% CI −0.08 to 0.05).

#### Associations With Sleep

In HCs, higher N3 SWD was associated with a lower DTI-ALPS index (r = −0.43, 95% CI −0.72 to −0.03); no sleep measure was associated with DTI-ALPS in PwMS (**Table 2**). In PwMS, stronger gBOLD–CSF coupling was associated with shorter TST (r = 0.49, 95% CI 0.05 to 0.77), and a higher arousal index (r = −0.48, 95% CI −0.77 to −0.04). Coupling was not associated with N3 microstructure in either group or with any sleep measure in HCs.

#### Associations With Clinical and Structural Measures

In PwMS, higher EDSS tended to be associated with a lower DTI-ALPS index (r = −0.44, 95% CI −0.74 to 0.01), with a similar estimate after adjustment for MD (r = −0.43, 95% CI −0.74 to 0.03). The association between coupling and EDSS was weak and imprecise (r = 0.25, 95% CI −0.22 to 0.62). Neither proxy was associated with disease duration (DTI-ALPS: r = 0.23, 95% CI −0.23 to 0.61; coupling: r = −0.22, 95% CI −0.60 to 0.25), lesion volume (DTI-ALPS: r = −0.22, 95% CI −0.60 to 0.25; coupling: r = 0.18, 95% CI −0.28 to 0.58), or ChP volume. Larger LV volume tended to be associated with weaker coupling (r = 0.44, 95% CI −0.00 to 0.74) but not with DTI-ALPS (r = −0.26, 95% CI −0.63 to 0.21) (**Table 2**).

## Discussion

This study combined 2 glymphatic MRI-derived proxies with overnight PSG in PwMS and yielded 3 main findings. First, the lower DTI-ALPS index in PwMS was accompanied by increased WM diffusivity and attenuated after adjustment for it, indicating that disease-related microstructural damage contributes to this widely used measure. Second, although gBOLD–CSF coupling did not differ between groups, it was stronger in PwMS with shorter and more fragmented sleep, consistent with a compensatory increase in waking CSF dynamics when sleep is impaired.^35,36^ Third, neither proxy was clearly associated with disability, disease duration, lesion volume, or ChP volume; only borderline associations emerged between lower DTI-ALPS and higher EDSS and between weaker coupling and larger LV.

Together, these findings suggest that DTI-ALPS should be interpreted with caution in MS and that gBOLD–CSF coupling may reflect a dynamic, sleep-dependent component of glymphatic-related physiology that is largely independent of disease burden. Because sleep disturbance is often treatable, this dynamic component may be a potential target for intervention.

MD within the DTI-ALPS ROIs was higher in PwMS, consistent with previous studies^3,37^ and with the sensitivity of MD to demyelination, axonal loss, gliosis, and edema, which affect periventricular white matter even in the absence of visible lesions. Because the DTI-ALPS index is a ratio of directional diffusivities measured in the same voxels, any process that alters the magnitude or anisotropy of diffusion in these regions affects the index independently of perivascular fluid transport.

Adjusting for MD attenuated the group difference, in agreement with a longitudinal study in which lower DTI-ALPS values in RRMS were no longer significant after accounting for MD.^37^ Most MS studies of DTI-ALPS have not adjusted for local diffusivity, including the only previous study relating DTI-ALPS to sleep in MS, which reported a lower index in patients with subjective sleep disturbances.^9^ Their findings may therefore partly reflect WM damage rather than perivascular fluid transport. Moreover, the assumption of preserved fiber orientation around medullary veins on which the index relies is likely violated in MS, in which visible and subtle demyelination affects periventricular WM.

The inverse association between N3 SWD and DTI-ALPS in HCs is unexpected: unlike gBOLD–CSF coupling, which increases after sleep deprivation, ^35,36,38^ DTI-ALPS decreases and only partly recovers after recovery sleep,^38^ suggesting that deeper sleep should accompany a higher index. If the index instead reflects periventricular WM integrity, this association may indicate more efficient neuronal synchronization, leading to fewer but potentially more effective slow oscillations. Its absence in PwMS, together with their slightly higher SWD, could indicate a sustained compensatory drive in which slow-wave generation is persistently upregulated and no longer dynamically modulated by WM integrity/glymphatic outflow. ROI placement may also contribute, as a recent sleep–wakefulness DTI-ALPS study on a heathy population found increased DTI-ALPS values during sleep in only 3 of 5 correctly placed ROI pairs.^39^

In PwMS, the association between DTI-ALPS and EDSS was borderline. Previous studies are inconsistent, with some reporting lower DTI-ALPS with higher EDSS and others no association,^3,37,40–42^ and in a longitudinal cohort a trend-level association disappeared after controlling for MD.^37^ DTI-ALPS was also not associated with lesion volume. Evidence on this relationship is also mixed: lower DTI-ALPS values with greater lesion volume have been reported in cohorts with smaller lesion volumes^3,42^ and in large cohorts including progressive MS,^3,43^ whereas a study in a cohort with lesion volumes comparable to ours found no association, consistent with our results.^41^ Differences in disease stage, sample size, ROI placement and the locations of lesions may contribute to these discrepancies.

Consistent with previous reports in early MS,^22,41^ PwMS had larger ChP and LV volumes than HCs, yet neither was associated with DTI-ALPS, unlike in a larger cohort in which ChP enlargement correlated with a lower DTI-ALPS index.^43^ This suggests that ChP enlargement may not necessarily induce perivascular changes detectable by this index. LV enlargement in MS mainly reflects atrophy,^23^ so the absence of an association with LV volume suggests that DTI-ALPS is not closely related to global structural disease burden. Atrophy is also thought to be accompanied by perivascular space enlargement,^6^ which would be expected to affect a perivascular diffusion measure; that DTI-ALPS was unrelated to LV volume is therefore more consistent with an index dominated by WM microstructure integrity than by perivascular fluid.

To date, several methodological limitations have been raised regarding DTI-ALPS: The index samples small periventricular WM regions, whereas tracer studies show prominent glymphatic enhancement in cortical regions;^10^ its initial correspondence with intrathecal gadolinium clearance^44^ has not been consistently replicated;^45^ perivascular spaces occupy only a small fraction of WM, making it unlikely that the index isolates perivascular diffusion from diffusion along fiber tracts; and results depend on ROI selection.^39^

In contrast to DTI-ALPS, gBOLD–CSF coupling was associated with objectively measured sleep in PwMS: coupling was stronger in those with shorter and more fragmented sleep. The shorter TST in PwMS resembles partial sleep restriction and waking gBOLD–CSF coupling increases after total or partial sleep deprivation.^35,36^ Stronger coupling in this context is unlikely to reflect better glymphatic function and more plausibly represents either a compensatory response to reduced sleep-dependent clearance ^35,36,46^ or a dysregulated, hypersynchronized state induced by sleep loss.^38^ These findings may appear to contradict those of Eide et al.,^47^ who showed with intrathecal tracer administration that a single night of total sleep deprivation reduced tracer clearance compared with normal sleep. However, a compensatory mechanism may exist yet be insufficient to fully counteract the effects of total sleep deprivation. Moreover, this compensation appears to be age dependent: studies comparing middle-aged (40–50 years) and older (65–75 years) individuals with younger adults suggest that it occurs mainly in younger individuals.^35,36^ Its absence in the participants of Eide et al., whose mean age was 44.7 years, may therefore have contributed to the persistent tracer accumulation.

The association with sleep fragmentation supports the same interpretation, because arousals interrupt the consolidated sleep periods during which clearance is thought to be most efficient. The absence of comparable associations in HCs suggests that physiological variation in sleep does not substantially alter CSF dynamics when baseline regulation is preserved, and the lack of association with N3 microstructure suggests that sleep quantity and continuity matter more for waking CSF dynamics than specific features of slow-wave sleep. Whether such compensation is sustainable over years of disturbed sleep, or eventually fails as disease progresses, is a clinically important question that only longitudinal studies can answer.

Coupling was not clearly associated with EDSS (r =0.25, 95% CI −0.22 to 0.62). The CI does not exclude a moderate association, and the narrow range of disability in this early MS cohort (EDSS 0–3.0) limited the ability to detect clinical correlations. A dynamic PET study of CSF clearance in MS likewise found no association with EDSS^48^.

Coupling was also unrelated to lesion volume. Global neural and CSF dynamics are unlikely to be driven by focal damage per se, and total lesion volume does not capture lesion heterogeneity, such as chronic active or remyelinated lesions, or the location of lesions relative to key CSF pathways. Coupling was likewise unrelated to ChP volume; although ChP enlargement in MS has been linked to inflammation and altered CSF production,^49^ gBOLD–CSF coupling reflects moment-to-moment CSF flow driven by neural and vascular activity rather than baseline CSF production. Dysregulation of CSF production alone may therefore not directly translate into changes in temporal coupling. Weaker coupling tended to accompany larger LV volume. Cerebral blood volume fluctuations have been proposed to exert mechanical forces on the ventricular walls that drive CSF flow,^19^ and gBOLD fluctuations correlate with short-term changes in LV volume.^25^ Ventricular enlargement could therefore attenuate the transmission of hemodynamic oscillations to CSF flow, for example by reducing the effective forces that move CSF out of the ventricles and disrupting the phase relationship between gBOLD and CSF signals. Alternatively, ventricular enlargement and decoupling could both reflect a shared disease process, such as atrophy-related changes in brain compliance. Because the CI of this association bordered zero, it requires confirmation before mechanistic conclusions can be drawn.

These findings have practical implications. First, DTI-ALPS is increasingly used as a glymphatic outcome and proposed as a biomarker in MS because it can be computed from routine diffusion MRI.^3,11^ Our data indicate that it should be interpreted only alongside mean diffusivity and should not be used as a standalone marker of glymphatic function in clinical studies or trials, where changes in WM integrity could be mistaken for changes in glymphatic function. Second, gBOLD–CSF coupling can be derived from a short resting-state fMRI acquisition without contrast agents and was sensitive to objectively measured sleep in PwMS, making it a feasible candidate marker of CSF dynamics in MS; such studies should account for recent sleep, on which waking CSF dynamics appear to depend. Third, the link between shorter, more fragmented sleep and altered CSF dynamics in PwMS provides a mechanistic rationale for systematically assessing sleep in MS care, where sleep disturbances are common,^8^ and for testing whether treating them modifies glymphatic-related dynamics and, ultimately, neuroinflammatory and clinical outcomes.

The absence of a group difference in gBOLD–CSF coupling, which to our knowledge has not previously been assessed in MS, has several possible explanations. Glymphatic dysfunction may not yet be pronounced enough in early, mildly disabled RRMS to alter this measure; the sample size limited power to detect subtle differences; and, if waking coupling increases to compensate for disturbed sleep, as our sleep associations suggest, compensation in some PwMS may have offset reduced coupling in others, masking a group effect. Assessment during nighttime sleep, when glymphatic activity is thought to be highest, and in patients with more advanced disease may reveal differences not apparent during wakefulness.

This study has limitations. The modest sample of PwMS with early, mildly disabled RRMS receiving various disease-modifying therapies limits precision and generalizability to progressive or untreated MS. Association analyses were exploratory and uncorrected for multiplicity, and several CIs were wide or bordered zero. gBOLD–CSF coupling was measured during wakefulness and compensatory waking mechanisms may have masked group differences that could emerge during sleep. Manual ROI placement could not be fully blinded because lesions are visible, although ROIs were placed according to predefined anatomical criteria. Sleep was recorded on a single night, which does not capture habitual sleep and may be affected by first-night effects. Finally, the cross-sectional design precludes causal inference. Longitudinal studies in larger cohorts spanning relapsing and progressive MS, with repeated sleep recordings and imaging during sleep, are needed to determine whether altered coupling precedes or predicts changes in sleep, disability, and brain structure.

In conclusion, multimodal glymphatic imaging in early RRMS shows that the DTI-ALPS index is confounded by WM damage, whereas gBOLD–CSF coupling tracks objectively measured sleep loss and fragmentation. gBOLD–CSF coupling thus emerges as a practical glymphatic-related marker in MS, and its sensitivity to sleep identifies sleep as a potential target to support brain clearance in people with MS.

## Acknowledgment

The authors thank all the individuals who participated in this study.

