## Supplementary material for "MRI-Based Glymphatic Markers and Their Associations with Polysomnographic Sleep in Relapsing-Remitting Multiple Sclerosis"

*Corresponding author: Mélanie Strauss

### **Supplementary Methods**

#### **MRI Acquisition**

All images were acquired on a 3T SIGNA PET/MR scanner (GE Healthcare, Chicago, IL) between 9 and 11 AM. The protocol included:

(1) Sagittal 3D fluid-attenuated inversion recovery (FLAIR): voxel size = 1 × 1 × 1.4 mm; 120 slices; matrix size= 256 × 256; repetition time (TR) = 7,200 ms; echo time (TE) = 118–120 ms and inversion time (TI) = 2,030–2,045 ms (minor intersubject variability due to vendor-prescribed automated protocol adjustments); echo train length = 180.

(2) 3D T1-weighted inversion-recovery spoiled gradient-recalled echo (GE BRAVO): voxel size = 1 × 1 × 1 mm; 176 slices; matrix size = 256 × 256; TR = 8.2 ms; TE = 3.1 ms; TI = 450 ms; flip angle = 12°.

(3) Axial pulsed-gradient spin-echo single-shot diffusion-weighted echo-planar imaging with b = 1000 s/mm², acquired along 32 noncollinear directions, plus 1 b = 0 image; voxel size = 2 × 2 × 2 mm; 70 slices; matrix size = 120 × 120; TR = 7,000 ms; TE = 70–85 ms; inversion time = 108 ms. An additional reversed phase-encoding b = 0 image was acquired for susceptibility distortion correction.

(4) Resting-state 2D gradient-echo echo-planar imaging: TR = 3,000 ms; TE = 34 ms; flip angle = 90°; field of view = 28 × 28 cm; matrix size = 96 × 96; slice thickness = 3 mm; frequency-encoding direction right-to-left; interleaved bottom-up slice acquisition. Four initial dummy volumes were discarded by the scanner; the 6-minute run yielded 120 volumes.

(5) 3D susceptibility-weighted imaging (SWI): voxel size = 0.8 × 0.8 × 1 mm; 156 slices; matrix size = 256 × 256; TR = 39–55 ms; TE = 20–25 ms; flip angle = 10°. Magnitude and phase images were reconstructed.

#### **Lesion Segmentation and Volumetry**

T2-FLAIR hyperintense lesions were segmented with LST-AI^e1^ in SPM12,^e2^ verified by an experienced MS neurologist (S.E.), and manually corrected when needed in ITK-SNAP (version 4.4.0).^e3^ Lesion filling was performed on T1-weighted and FLAIR images with FSL.^e4^

Lateral ventricle volumes were obtained by parcellating the lesion-filled T1-weighted image^e5^ with FreeSurfer (version 6.0.1)^e6^ and the Desikan-Killiany atlas^e7^ in FSL (version 6.0.7.6)^e8^ and were normalized to estimated total intracranial volume (eTIV). Choroid plexus volumes were obtained with the method of Tadayon et al.,^e9^ because the FreeSurfer segmentation of this structure has limited accuracy.^e9–e11^ All segmentations were reviewed by a neurologist (S.E.) blinded to clinical data before normalization to eTIV.

#### **DWI Preprocessing and DTI-ALPS Index**

Diffusion-weighted images were preprocessed with a modified version of the pipeline of Liu et al. (github.com/gbarisano/alps).^e12^ Preprocessing included denoising and Gibbs-ringing removal with MRtrix3,^e13^ susceptibility-induced distortion correction with FSL TOPUP,^e14^ and eddy-current and motion correction with FSL EDDY.^e15^ The preprocessed data were fitted with FSL dtifit to obtain mean diffusivity (MD) and fractional anisotropy maps and tensor images, from which diffusivity maps along the x (Dxx), y (Dyy), and z (Dzz) axes were extracted.

For each participant, the lesion mask and SWI images were coregistered to DWI space with ANTs (github.com/antsx/ants), and all registrations were visually inspected.

#### **Resting-State fMRI Preprocessing and Signal Extraction**

Functional data were corrected for slice timing and then for motion. Temporal band-pass filtering (approximately 0.01–0.1 Hz) and linear detrending were applied, without spatial smoothing. Denoised, motion-corrected data were used for gBOLD extraction. CSF signal extraction was performed on data that underwent slice-timing correction and temporal filtering but no motion correction, to preserve the slice-position information required for CSF inflow analysis.

For the gBOLD signal, the Harvard-Oxford cortical structural atlas (RRID:SCR_001476)^e7,e16–e18^ was nonlinearly registered to T1 space, binarized, intersected with the binary gray matter mask obtained from T1 segmentation with CAT12 (neuro-jena.github.io/cat),^e19^ and linearly transformed to fMRI space. Subcortical regions were excluded because of their higher susceptibility to vascular artifacts. Time series were z-normalized and averaged across all voxels of this mask, yielding a gBOLD signal that reflects both the amplitude and the synchronization of global brain activity.^e20^

The CSF inflow signal was extracted from the lowest slice of the acquisition, where sensitivity to the inflow effect is maximal: fresh spins entering the imaging volume, unaffected by previous radiofrequency pulses, increase the fMRI signal.^e21^ The CSF mask was obtained by segmenting the fMRI data with the unified segmentation approach^e22^ in SPM12 and thresholding at a probability of 1.00 to retain only the highest-probability CSF voxels. The CSF signal was expressed as percentage change from its temporal mean.

To test whether the group-averaged coupling exceeded chance, a null distribution was generated by pairing gBOLD and CSF signals from different participants (10,000 permutations), and the observed group-averaged cross-correlation was compared with this distribution.

Consistent with previous studies of gBOLD–CSF coupling,^e20,e21,e23–e26^ we did not regress out the global or CSF signals, which were the signals of interest. Although head motion has been proposed as a source of resting-state fMRI signal changes, recent findings suggest that it more likely reflects arousal-related global BOLD fluctuations than drives them;^e27^ motion parameter regression was therefore not applied, to avoid attenuating the global BOLD signal.

#### **SWI Preprocessing**

SWI images were reconstructed from T2*-weighted magnitude and phase images. Phase data were normalized, converted to radians (mapped linearly to the range −π to π), and high-pass filtered to remove low-frequency background contributions. A phase mask was generated to enhance susceptibility contrast, particularly of veins: voxels with nonnegative phase were assigned a weight of 1, whereas negative phase values were scaled linearly between 0 and 1, with greater attenuation toward −π.^e28^ The final SWI image was obtained by multiplying the phase mask with the magnitude image.

#### **Statistical Analysis Details**

Partial correlations were computed by regressing the covariates (age and sex, and MD where stated) out of both variables and correlating the residuals; P values (used only internally) and 95% CIs account for the covariates (t test with n − 2 − k degrees of freedom; Fisher z standard error 1/√(n − 3 − k), where k is the number of covariates). Group differences were estimated by ordinary least squares with group, age, and sex as predictors; the coefficient for group (PwMS minus HCs) and its 95% CI are reported.

**Supplementary Table S1** Disease-Modifying Therapies in Patients

| **Disease-modifying therapy** | **PwMS (n = 22), n (%)** |
| --- | --- |
| Teriflunomide | 2 (9.1) |
| Dimethyl fumarate | 1 (4.5) |
| S1P receptor modulators (fingolimod, ozanimod) | 4 (18.2) |
| Cladribine | 1 (4.5) |
| Natalizumab | 6 (27.3) |
| Anti-CD20 therapies (ocrelizumab, ofatumumab) | 8 (36.4) |

Abbreviations: PwMS = patients with relapsing-remitting multiple sclerosis; S1P = sphingosine-1-phosphate.
